# ENLIGHT Consensus Checklist and Guidelines for reporting laboratory studies on the non-visual effects of light in humans

**DOI:** 10.1101/2023.03.17.532785

**Authors:** Manuel Spitschan, Laura Kervezee, Renske Lok, Elise McGlashan, Raymond P. Najjar, the ENLIGHT Consortium

## Abstract

Beyond vision, light has wide-reaching effects on human health and well-being. However, there is no consensus on reporting light characteristics in studies investigating non-visual responses to light. This project aimed at developing a reporting checklist for laboratory-based investigations on the impact of light on non-visual physiology. To this end, a four-step modified Delphi process (three questionnaire-based feedback rounds and one face-to-face group discussion) involving international experts was conducted. Across these four rounds, an initial list of 61 items related to reporting light-based interventions was condensed to a final checklist containing 25 items, based upon consensus among experts (final n=60). Nine of these items were determined to be necessary to report regardless of the specific research question or context. A description of each item was provided in the accompanying guidelines. Most participants (92%) reported being satisfied or very satisfied with the consensus process, checklist, and guidelines. The ENLIGHT Checklist and Guidelines are the first consensus-based guidelines for documenting and reporting ocular light-based interventions for human studies. The implementation of the checklist and guidelines will enhance the impact of light-based research by ensuring comprehensive documentation and reproducibility and enabling data aggregation across studies.

## Introduction

Light exerts powerful effects on our physiology and behavior beyond enabling vision (1, 2). One of the primary non-visual functions of light is the synchronization of the circadian clock (3). The adaptability of the circadian clock allows us to adapt to seasonal changes in day length, travel across time zones, and changes in our sleep-wake schedule. However, abnormal shifts in circadian timing can be associated with pathology, including a number of mood and sleep disorders (4–6). The circadian clock is implicated in nearly all disease states, including metabolic conditions, cardiovascular health, and some cancers (7–10). Critically, the timing of the circadian clock can impact the efficacy of treatment outcomes (11), and therefore understanding circadian timing and the impact that an individual’s light exposure may have on their clock could profoundly impact treatment outcomes. In addition to effects on the circadian clock, light has several direct effects on functioning, including increasing alertness (12, 13), improving mood (2), and causing changes in cognitive brain function (14).

The non-visual effects of light are mediated by a population of cells in the retina called intrinsically photosensitive retinal ganglion cells (ipRGCs) (15–18). These cells integrate signals from traditional photoreceptors (rods and cones) and project from the retina to the circadian master clock in the suprachiasmatic nucleus (SCN) (19), as well as several other brain areas involved in mood, alertness, and pupillary responses. ipRGCs are predominantly sensitive to short-wavelength (blue) light due to the light-sensitive protein called melanopsin (16, 20–23). Therefore, the spectral quality of a light source determines its impact on non-visual functions, in addition to the intensity, timing, and temporal pattern of light exposure.

The non-visual effects of light vary significantly among individuals (24–26). Furthermore, non-visual responses can be elicited by very low light levels and very rapidly (27–29). Even minor differences in the intensity, pattern, spectral quality, or timing of light stimuli in clinical and basic research studies may result in substantial differences in the response observed. Therefore, it is becoming increasingly necessary that measurement and reporting practices in lighting research are standardized. Standardizing light reporting also allows for direct comparisons between studies, enabling meta-analyses to be conducted more effectively. There have been several proposals for common reporting schemes (30–33), the oldest dating back to 1991 (32). As neurobiological studies on the non-visual effects of light are highly resource-intensive, often taking multiple years to complete, a consensus checklist represents a significant step forward for the field.

This study aimed to produce a specific reporting checklist and accompanying guidelines for light and study characteristics in laboratory-based, light-centered interventions conducted in human research participants to study the effects of ocular light exposure on non-visual physiology. The ENLIGHT (**E**xpert **N**etwork on **LIGHT** Interventions: **ENLIGHT**) checklist is intended to assist reviewers, editors, and readers in appraising the completeness and applicability of the findings from a study, as well as enable the synthesis of data across published work. Researchers may also use the checklist to evaluate and organize the information critical to light studies when designing a study or preparing a grant application.

## Results

### Novel consensus checklist and guidelines for light-based interventions

The ENLIGHT Checklist and Guidelines were developed using a four-step modified Delphi process (34). This consisted of preliminary work to identify an appropriate list of initial items for evaluation (see **Methods** and **Supplementary Table S2**), three survey-based rounds (Round 1, 2 and 4), and one round of face-to-face discussion (Round 3). The goal of Round 1 was to assess ratings of the importance of the list of items identified in the preliminary work and the initial drafting of the checklist. In Round 2, experts were asked to evaluate the initial draft of the checklist, and to provide input on the preferred format of items. Round 3 consisted of face-to-face discussions with experts to clarify questions or concerns from participants, discuss the scope of the guidelines accompanying the checklist, and discuss how to maximize the impact and adoption of the ENLIGHT Checklist and Guidelines. In Round 4, the final version of the checklist and accompanying guidelines were reviewed, and experts were asked to indicate which items should be mandatory. The final ENLIGHT Checklist is a convenient, fillable form-based PDF.

### Round 1: Importance of preliminary items, gathering of additional items, and initial checklist drafting

Of the 115 invited experts, 65 participants completed the first survey (see **Figure 1** for a flow chart of participant recruitment and **Table 1** for demographic information). Participants rated the importance of 61 items and quantities for reporting on a scale from 1 (very unimportant) to 7 (very important) across 12 domains (**Figure 2**). Twenty-four items reached the threshold for definite inclusion, i.e., these were rated with a score of at least “5 – important” by ≥75% of participants. The 37 remaining items were rated as either “unimportant” (a score of less than 3) or “unknown”, with the exact per-item percentage ranging between 8 and 60%. None of the items was rated “unimportant” by ≥75% of experts and therefore none met the threshold for definite exclusion.

**Figure 1:**
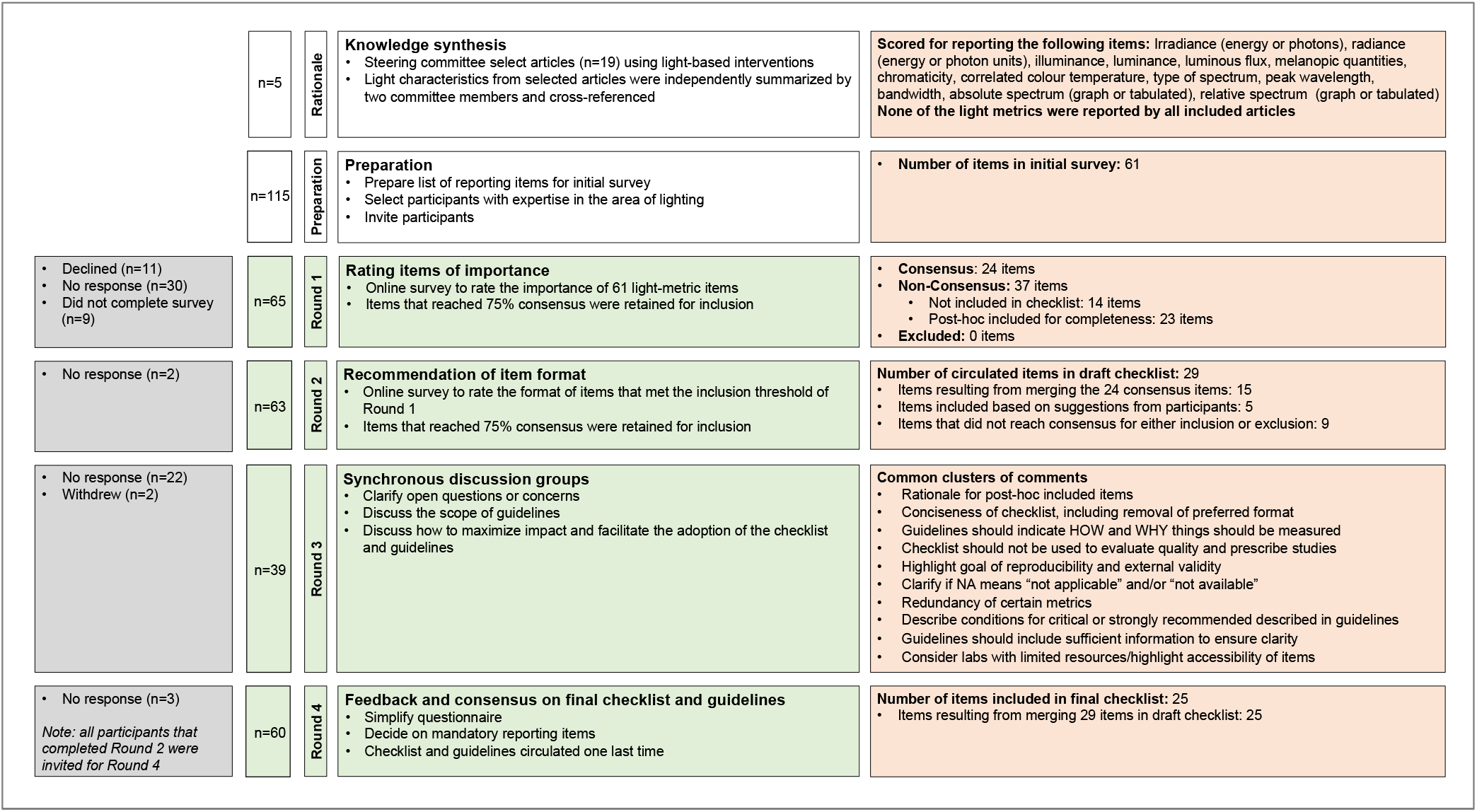
General overview of participant inclusion, the consensus rounds, and item selection for the checklist.

**Figure 2:**
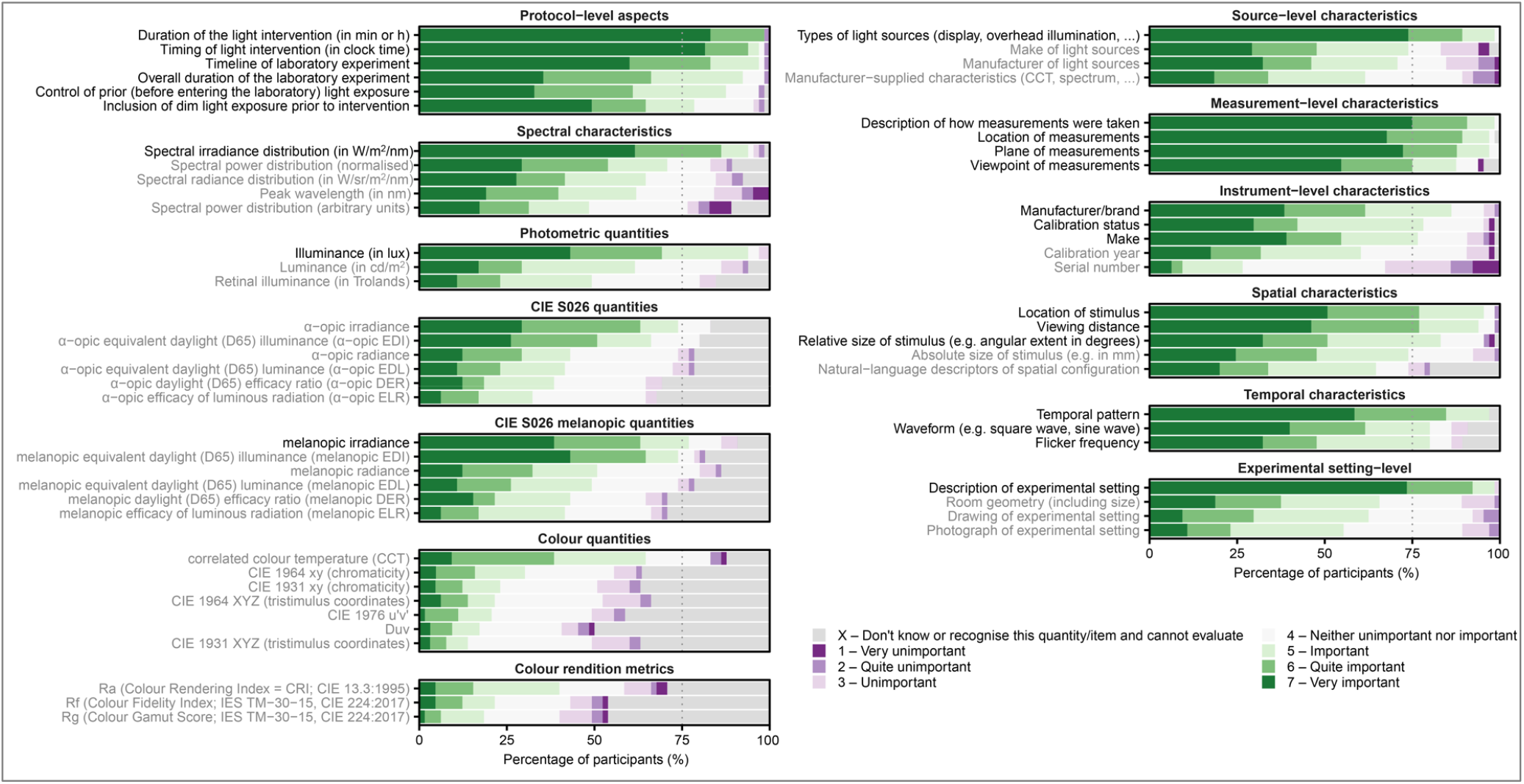
Ratings of initial list of 61 potential checklist items as unimportant–important by participants in Round 1 (n=65 participants). Items in black: items that reached the consensus threshold for inclusion. Items in gray: items that did not reach the threshold for either inclusion or exclusion.

**Table 1.**
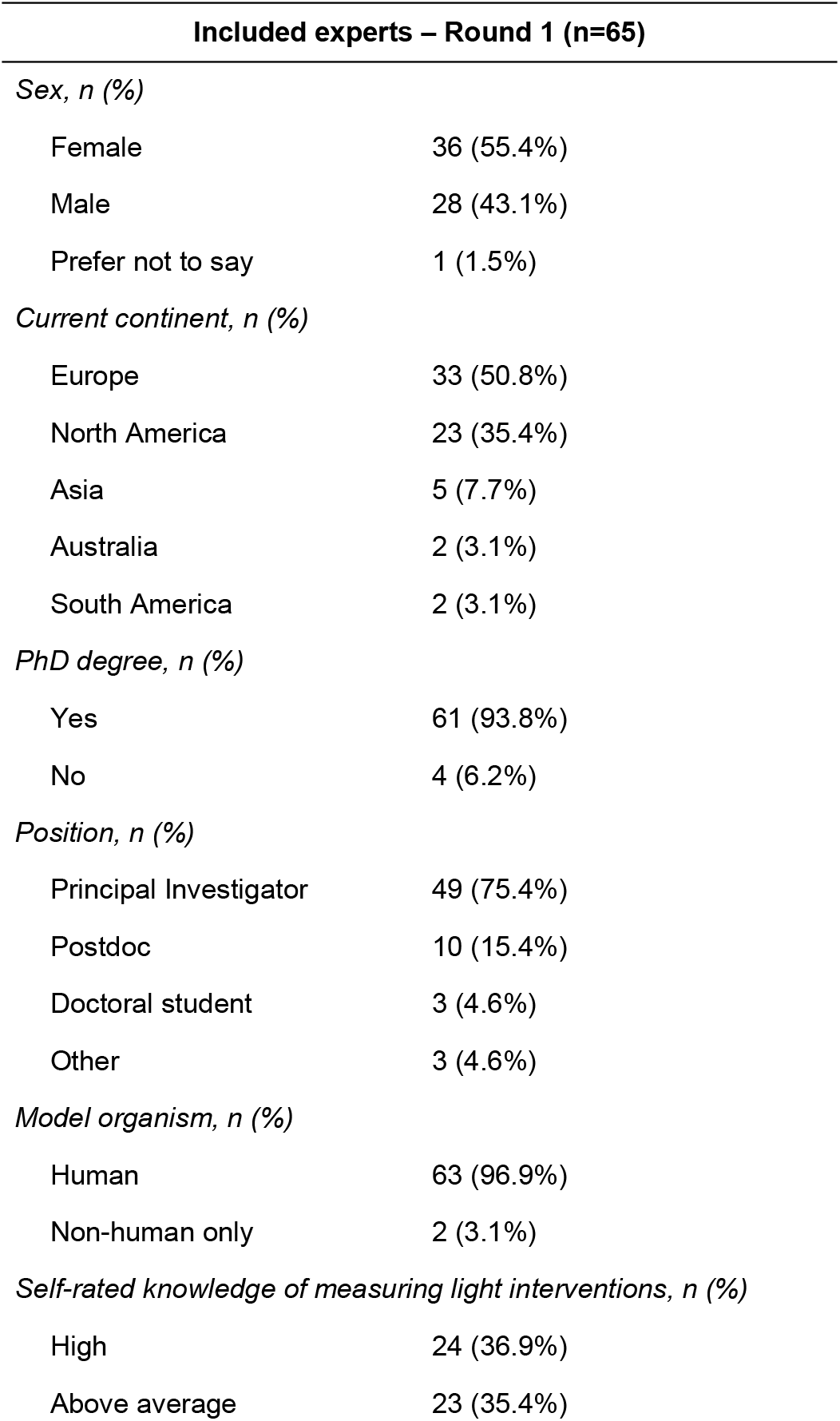
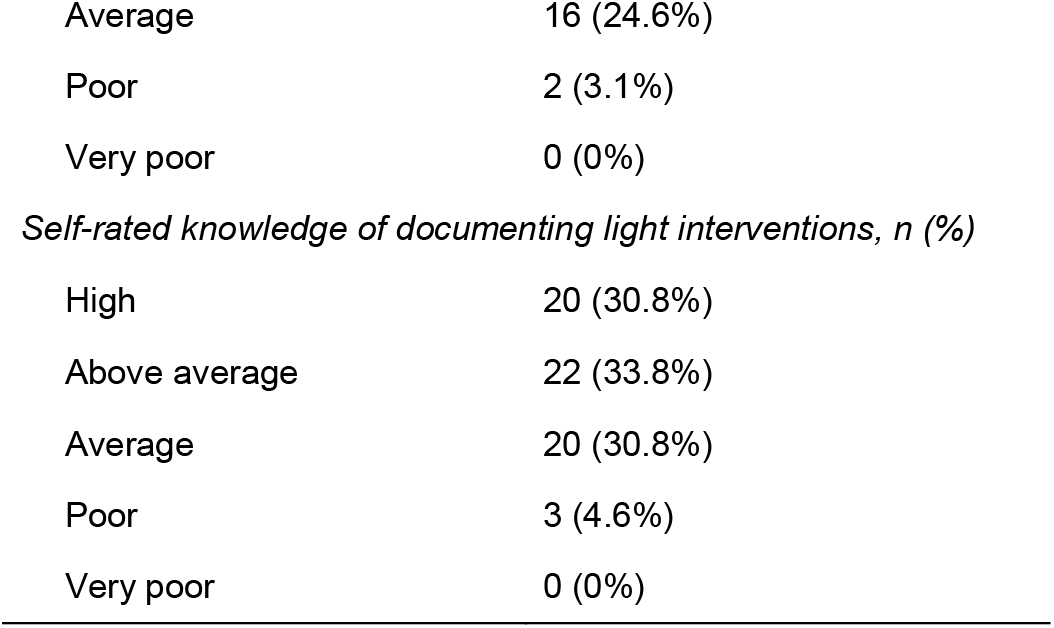
Demographic characteristics of participants of Round 1 (n=65)

| **Included experts – Round 1 (n=65)** |  |
| --- | --- |
| *Sex, n (%)* |  |
| Female | 36 (55.4%) |
| Male | 28 (43.1%) |
| Prefer not to say | 1 (1.5%) |
| *Current continent, n (%)* |  |
| Europe | 33 (50.8%) |
| North America | 23 (35.4%) |
| Asia | 5 (7.7%) |
| Australia | 2 (3.1%) |
| South America | 2 (3.1%) |
| *PhD degree, n (%)* |  |
| Yes | 61 (93.8%) |
| No | 4 (6.2%) |
| *Position, n (%)* |  |
| Principal Investigator | 49 (75.4%) |
| Postdoc | 10 (15.4%) |
| Doctoral student | 3 (4.6%) |
| Other | 3 (4.6%) |
| *Model organism, n (%)* |  |
| Human | 63 (96.9%) |
| Non-human only | 2 (3.1%) |
| *Self-rated knowledge of measuring light interventions, n (%)* |  |
| High | 24 (36.9%) |
| Above average | 23 (35.4%) |
| Average | 16 (24.6%) |
| Poor | 2 (3.1%) |
| Very poor | 0 (0%) |
| *Self-rated knowledge of documenting light interventions, n (%)* |  |
| High | 20 (30.8%) |
| Above average | 22 (33.8%) |
| Average | 20 (30.8%) |
| Poor | 3 (4.6%) |
| Very poor | 0 (0%) |

In Round 1, participants also had the opportunity to suggest additional items for the checklist and provide open-ended feedback. Three clusters of common feedback were identified:

- **Suggestions for additional checklist items.** These items were mostly related to observer-level characteristics specific to light interventions (e.g., pupil dilation, ocular functioning, and timing of light interventions relative to individual circadian or sleep time).
- **Scope of checklist.** The experts noted that the importance and relevance of items highly depend on the specific study.
- **Organization of the checklist**. The experts noted that the checklist should be concise, simple to use and not add additional burden on researchers for documenting their research.

Based on the last concern, it was decided to condense and combine some of the 24 items that reached consensus for inclusion, resulting in 15 items. Subsequently, a draft checklist was created that contained 29 items. This included the 15 condensed items that reached consensus and five new items based on suggestions from participants. After internal discussions among the ENLIGHT Steering Committee, an additional nine items that did not reach consensus for *either* inclusion or exclusion were also included in the 29-item checklist to ensure the checklist covered all major aspects of lighting, as for some aspects, no individual items reached consensus at this stage.

### Round 2: Draft checklist evaluation and format specification

Sixty-three participants (97% of participants that completed Round 1) completed Round 2. No items on the initial draft of the checklist were rated by the majority as “should not have been included”, and no items from Round 1 that were excluded from the initial draft were rated as “should have been included” by the majority of experts (**Supplemental Figure S1**), suggesting that there was support for the initial draft. Consequently, all 29 items were retained for discussion in Round 3.

In addition, participants were asked to indicate their preference for the reporting format of the different items in the draft checklist. For most items, text was the preferred format, although for some items a figure or table was preferred by the majority of participants (**Supplemental Figure S2**). For example, “figure” was the preferred format to report the timeline of the experiment, while “table” was preferred to report α-opic (ir)radiances.

### Round 3: Synchronous discussions

The 63 participants that completed Round 2 were invited to participate in the synchronous discussions (Round 3 of the modified Delphi process). Participants who agreed to participate (n=41) were split into six working groups that ranged in size from 4 to 11 participants, with a median of 7 participants per group. One video call was held for each group. Two members of the ENLIGHT Steering Committee (one moderator and one note-taker) participated in each call. Overall, 39 out of 41 participants attended their respective video calls. Two participants withdrew after being allocated to a group. The synchronous discussions focused on three objectives: (1) To clarify any open questions or concerns from participants; (2) To discuss the scope of the guidelines accompanying the checklist; and (3) To discuss how to maximize impact and facilitate the adoption of the ENLIGHT Checklist and Guidelines. Two Steering Committee members reviewed video call minutes (R.L., L.K.). Recurrent comments and discussion points raised by participants were identified, discussed by the ENLIGHT Steering Committee, distilled, and grouped under ten common themes raised by multiple participants, within the three objectives of this round (**Supplementary Table S3**). These themes were used to simplify, condense, and improve the checklist as well as to inform the preparation of the guidelines. Feedback on actions taken by the Steering Committee was provided to the ENLIGHT Consortium participants in Round 4 of the Delphi Process.

Following Round 3 of the Delphi process, the 29 items in the initial draft checklist were condensed to 25 items based on the feedback to further simplify and shorten the checklist. Furthermore, the wording of nine checklist items was improved. For example, “flicker frequency” was replaced by “flash frequency” to clarify that this item pertains to the intentional temporal pattern of the light stimulus. Likewise, the item ‘pupil dilation’ was reworded to “pupil size and/or dilation” to clarify that this item relates to both the description of any methods used to pharmacologically dilate the pupil as well as any methods used to measure and/or control for pupil size. In addition, based on specific comments by participants, general textual/structural improvements were made in the checklist, including (1) the removal of text/table/figure designation; (2) the specification that all light sources used should be reported; and (3) replacement of the term “light intensity” to “light level” for accuracy.

### Round 4: Finalization of checklist and guidelines

In Round 4, all participants that had completed Round 2 were asked to vote on which items on the checklist they deemed essential to be reported regardless of experimental context and to provide qualitative feedback on the final checklist and accompanying guidelines. In total, 60 participants completed this round. Nine items reached consensus on being essential, i.e., these items were rated by more than 75% of the participants as essential (**Supplemental Figure S3**). Based on open-ended feedback, minor textual changes were made to the checklist for accuracy. These changes included renaming the items ‘color quantities’ and ‘color rendition metrics’ to ‘color appearance quantities’ and ‘color rendering metrics’, respectively. Following feedback from the participants, some references in the accompanying guidelines were either removed or replaced with better-suiting ones. Lastly, 92% of participants reported being satisfied or very satisfied with the consensus process as well as the final checklist and guidelines (**Figure 3**).

**Figure 3:**
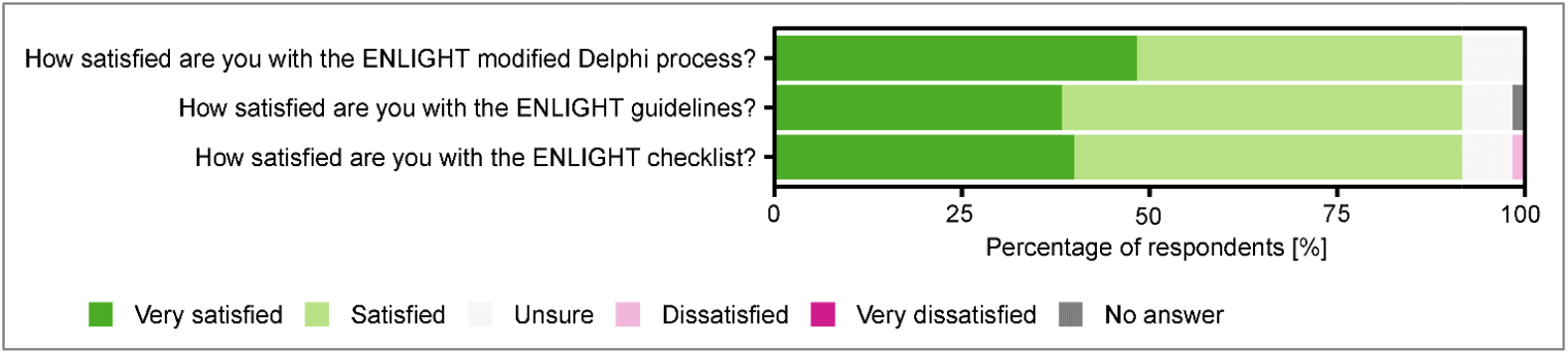
Ratings of participant satisfaction with the ENLIGHT process, guidelines, a**nd** final checklist (n=60 participants in Round 4).

A list of items included in the final checklist and a short description of each item is presented in **Table 2**. A fillable checklist is available as **Supplemental File S1**. The final guidelines are available as **Supplemental File S2**.

**Table 2.** Items in final ENLIGHT checklist.

| Item | Description |
| --- | --- |
| **A. Study characteristics** |  |
| *A.1. Protocol-level characteristics* |  |
| Description of experimental setting* | Describe the experimental setting (e.g., room geometry). |
| Timeline of experiment (including timing and duration of light)* | Provide an overview of the timing of key study events, including timing and duration of light exposure. |
| Pre-laboratory sleep-wake/rest-activity behaviour | Describe the pre-laboratory sleep-wake or rest-activity behaviour (e.g., any measurement of participants' sleep-wake or rest-activity behaviour prior to entering the laboratory). |
| Pre-laboratory light exposure | Describe the pre-laboratory light exposure, including whether participants were given any instructions related to light exposure. |
| Immediate prior light exposure (in laboratory) | Describe the in-laboratory light conditions immediately prior to the experimental light exposure. |
| *A.2. Measurement-level characteristics* |  |
| Measurement plane (e.g., horizontal or vertical)* | Describe the plane in which light measurement(s) were performed. |
| Measurement viewpoint and location* | Describe the location and direction at which the light sensor was placed during light measurements. |
| Type, make and manufacturer of the measurement instrument* | Describe the instrument being used to take each light measurement, including the manufacturer, type, make and model of the device. |
| Calibration status of the instrument | Describe the calibration status of the light sensor that was used to take each light measurement. |
| *A.3. Participant-level characteristics* |  |
| Ocular health and functioning* | Provide any details on health and functioning of the participants' eyes. |
| Pupil size and/or dilation | Describe any pupil size measurements and/or whether pupils were pharmacologically dilated during the experimental protocol. |
| Relative time (e.g. to circadian phase or sleep) | Describe the time of the experimental light exposure relative to the participants' sleep or circadian timing. |
| **B. Light characteristics** |  |
| *B.1. Light sources* |  |
| Light source type(s)* | Tick all relevant boxes to indicate the type(s) of background and experimental light sources used in the study. |
| Type, make and manufacturer of the light source* | Describe the type, make, and manufacturer of the light source(s) used in the study. |
| Use of wearable filtering apparatus (e.g., blue-blocking glasses) | Describe any wearable device(s) that modifies the absolute flux level or relative spectral distribution, or both, of light passing through it. |
| *B.2. Light level characteristics* |  |
| Illuminance (lux) and/or luminance (cd/m<sup>2</sup>)*# | Provide the illuminance and/or luminance of the experimental light condition(s) used in the study. |
| Spectral irradiance and/or radiance distribution<sup>#</sup> | Provide the spectral irradiance and/or radiance distribution of the experimental light condition(s) used in the study. |
| α-opic irradiance and/or radiance (including melanopic)<sup>#</sup> | Provide the α-opic irradiance and/or radiance of the experimental light condition(s) used in the study. |
| α-opic equivalent daylight illuminance and/or luminance (EDI/EDL, including melanopic)<sup>#</sup> | Provide the α-opic equivalent daylight illuminance and/or luminance of the experimental light condition(s) used in the study. |
| *B.3. Colour characteristics* |  |
| Peak wavelength and bandwidth | Provide the peak wavelength and bandwidth of the experimental light condition(s). Note that these metrics are most relevant for monochromatic or narrowband light sources. |
| Colour appearance quantities (any) | Provide colour appearance quantities of the experimental light condition(s), such as any metric describing position in a chromaticity diagram or color space, or correlated colour temperature, CCT (Tc). |
| Colour rendering metrics (any) | Provide any colour rendering metrics, such as the Colour Fidelity Index, Rf. |
| **B.4. Temporal and spatial characteristics** |  |
| Location of stimulus and viewing distance* | Describe the location of the light stimulus relative to the participant, and/or the relative distance between the light stimulus and the participant. |
| Temporal pattern (including flash frequency and waveform) | Describe the temporal pattern of the light sequence (e.g., the flash frequency or inter-stimulus interval) and the waveform (e.g., square, sinusoidal). |
| Relative or absolute size of the stimulus | Describe the size of the light stimulus, either absolute or relative (in relation to the visual field). |

\*Items marked with an asterisks reached consensus for being essential to report in any study regardless of experimental context.
#Luminance and radiance metrics (as opposed to illuminance and irradiance) are mainly relevant for emissive surfaces.

## Discussion

The ENLIGHT Checklist and Guidelines have been developed to provide guidance on the reporting of human laboratory studies deploying ocular light interventions. A modified four-step Delphi process was implemented, involving three questionnaire-based rounds and one round of face-to-face discussions. An initial list of 61 items was condensed into a final checklist of 25 items, 9 of which were determined by experts to be necessary to report regardless of the specific research question or context.

The ENLIGHT Checklist is a form-based fillable PDF and is available to download in **Supplemental File S1**. Authors are encouraged to fill in the checklist prior to the submission of a manuscript to ensure completeness in reporting. The checklist is not intended to serve as a guide to evaluating design quality, only to facilitate complete reporting of relevant study and light characteristics and enhance reproducibility and external validity. The ENLIGHT Guidelines for completing the checklist are included in **Supplemental File S2**. The guidelines provide definitions and examples for items included in the checklist, along with additional resources and tools for calculating or understanding metrics referenced in the checklist. Many of the items included in the final version of the checklist do not require any special measurement equipment and can be documented and reported at no cost. Therefore, uptake of the ENLIGHT Checklist and Guidelines can be readily achieved. Ongoing feedback and suggestions can be submitted online as an “issue” on our GitHub repository (https://github.com/ENLIGHT-Project/ENLIGHT-Guidelines-Checklist), which will be used to inform potential revisions of the checklist and guidelines in the future. Although the ENLIGHT Checklist is intended to be used for human laboratory studies, modifications could be made to apply the checklist and guidelines to other contexts, including field studies.

Achieving consensus within a community of researchers with different backgrounds, interests, expertise, and training necessarily leads to compromises. A balance must be achieved between adequate and complete reporting, and the sometimes extensive resources required for particular measurements or metrics to be taken. We, the ENLIGHT Steering Committee, believe the consensus reached shows an appropriate balance between these competing interests, given the current availability of resources and expertise.

We would like to note that one item which did not reach consensus but appeared multiple times in qualitative feedback from experts as an essential metric underpinning the reproducibility of studies. Spectral power distribution (in Section B.2. *“Light level characteristics”* of the checklist, **Table 2**) was rated by 73% of experts as essential to report in all contexts, falling just below the threshold for consensus (≥75%). Spectral measurements enable the calculation of many other metrics of interest, and therefore are the simplest way to support reproducibility and comparison between studies. We acknowledge that the measurement of spectral distribution requires specialized equipment, which may not be available to all researchers, but nonetheless encourage authors to report this item whenever it is available.

Given the complex nature of measuring and reporting light metrics in human studies, the committee highlights that researchers and practitioners wishing to employ light as interventions must be appropriately trained in optical radiation metrology. During the consensus process, some experts highlighted that there is a lack of field-specific, accessible educational materials. The level of training necessary to perform some of the measurements poses a significant barrier to measuring and reporting specific metrics. The accompanying ENLIGHT Guidelines provide some tools and resources for understanding and calculating the items covered by the checklist. However, increasing accessible and appropriate education tools or materials will aid in increasing the ENLIGHT Checklist usage.

The ENLIGHT Checklist and Guidelines are the first consensus-based guidelines for documenting and reporting light-based interventions for biomedical studies. The checklist and guidelines were derived through a systematic process involving in-depth interactions with experts in the field and future users of the checklist. Significant interindividual differences exist in non-visual responses to light, and effects can be seen with even very low-level exposure. Therefore, minor differences in the delivery method, intensity, or spectral composition of light exposure may result in substantial variation in responses observed. In conclusion, the ENLIGHT Checklist and Guidelines represent a crucial step in improving the documentation of research on the physiological and biobehavioral effects of light, making this work more reproducible and fit for large-scale data synthesis.

## Methods

### Ethical approval and registration

The ENLIGHT project received ethical approval from the University of Oxford Medical Sciences Interdivisional Research Ethics Committee (MS IDREC) (approval number R78618/RE001). All participants gave informed consent to be part of this study. The study was registered with the EQUATOR Network (https://www.equator-network.org/library/reporting-guidelines-under-development/reporting-guidelines-under-development-for-other-study-designs/#ENLIGHT), and the protocol was pre-registered on the Open Science Framework (https://doi.org/10.17605/OSF.IO/XR965).

### Code, materials and data availability

All code, materials and data are available on GitHub (https://github.com/ENLIGHT-Project/), containing the survey configurations used in JISC Online Surveys (https://github.com/ENLIGHT-Project/ENLIGHT-Survey), the data (https://github.com/ENLIGHT-Project/ENLIGHT-Data), and the archival guidelines (https://github.com/ENLIGHT-Project/ENLIGHT-Guidelines-Checklist). All data and materials are licensed under CC-BY-NC-ND, and all code is licensed under GPLv3.

### Protocol and ENLIGHT Steering Committee

The ENLIGHT Checklist and Guidelines were developed in accordance with the EQUATOR toolkit for developing a reporting guideline (https://www.equator-network.org/) through a modified Delphi consensus process that took place between December 2021 and December 2022 consisting of (1) a pre-round where the project team went through a literature review, identified a set of potential reporting-related items and a group of participants with an established track record in light-based interventions and (2) four feedback rounds (three questionnaire-based and one face to face discussion) detailed in the procedure section below.

A five-member steering committee was established to coordinate the development process of ENLIGHT, consisting of scientists with expertise in the visual and non-visual effects of light. The ENLIGHT Steering Committee consists of M.S., a visual neuroscientist with expertise in visual and circadian neuroscience, L.K., a chronobiologist with expertise in human physiology and neuroscience, R.L., a chronobiologist with expertise in light, circadian rhythms and alertness, E.M., a chronobiologist with expertise in light, circadian rhythms, and mental health, and R.P.N., a visual neuroscientist with expertise in circadian biology, light and ocular diseases. The ENLIGHT Steering Committee coordinated the modified Delphi consensus process, including selecting participants, designing and distributing the online surveys for the modified Delphi consensus process, organizing and moderating the face-to-face consensus meetings, analyzing data, and drafting the ENLIGHT Checklist and complementary guidelines. The ENLIGHT Steering Committee had regular online meetings and met in person in three one-week visits to coordinate and finalize the Delphi process as well as the ENLIGHT Checklist and Guidelines.

### Participants and consortium formation

Potential participants with experience in laboratory-based studies in human participants on the non-visual effects of light were invited to participate in this exercise through purposive sampling. To ensure the broadest representation of feedback, an effort was made to invite participants at different career stages, of any gender, from different geographical locations, and working at a variety of academic and industrial institutions. In the invitation email, participants were also asked to provide recommendations and contact details of additional suitable participants to be included in the exercise Participants who completed Rounds 1, 2 and 4 of the consensus process were invited to join the ENLIGHT Consortium and are acknowledged by name within a group authorship model (**Supplemental Table S1**). Participants were not remunerated for their participation.

### Literature review and rationale

To motivate the development of a reporting checklist and guidelines, we reviewed the metrics and concepts reported in human studies of the non-visual effects of ocular light exposure. We identified a recent paper that synthesized findings across studies of the non-visual effects of light to attempt to answer questions relating to the intensity and spectral composition of light (29). The references cited in this paper were examined for reported light characteristics. Two members of the ENLIGHT Steering Committee summarized these for reporting the following items: Irradiance (energy or photons), radiance (energy or photon units), illuminance, luminance, luminous flux, melanopic quantities, chromaticity, correlated color temperature, type of spectrum, peak wavelength, bandwidth, absolute spectrum (graph or tabulated), relative spectrum (graph or tabulated). Results were then cross-referenced between members. While all articles reported at least one measure of light intensity (either irradiance, radiance, illuminance, luminance, or luminance flux), no single measure of light intensity was reported across articles. None of the other metrics were reported by all 19 articles (see **Supplemental Table S2**).

### Procedure

#### Preliminary work: Selection of concepts

We identified concepts and items that could be used in the ENLIGHT Checklist. These were based on the steering committee’s domain knowledge, as well as by consulting the relevant literature. In identifying concepts, we consulted a number of key references. As early as 1991, Remé, Menozzi and Krueger (32) proposed a reporting scheme. Spitschan *et al.* (31) proposed a series of items for reporting interventions involving light in the field of chronobiology, sleep research, and environmental psychology. Knoop *et al.* (33) developed a workflow for identifying which quantities to measure and report in research on the non-visual effects of light. Finally, the International Commission on Illumination (CIE) released a technical note, CIE TN 011:2020, discussing items to report in studies on ipRGC-influenced responses to light (30).

Considering both specific metrics and study protocol aspects identified by these resources, as well as broad aspects of study design or lighting, which are consistent across the existing resources, an initial pool of 61 items was generated. These items covered protocol, experimental, measurement, instrument, and source level characteristics, as well as the spectral, photometric, color, spatial and temporal aspects of the light source.

#### Round 1: Importance of preliminary items, gathering of additional items, and initial draft of checklist

In Round 1, participants were invited by email to complete an online survey using a web-based survey tool hosted by the University of Oxford (JISC Online Surveys). After reading a participant information sheet, participants were asked to complete an informed consent form and then to rate the importance of a set of preliminary items identified in the *Pre-round* on a 1-7 scale, with the following options: 1 – Very unimportant, 2 – Quite unimportant, 3 – Unimportant, 4 – Neither unimportant nor important, 5 – Important, 6 – Quite important, 7 – Very important. To allow for participants being unable to assess the importance of a specific item, we also included a further open category (X – Don’t know or recognize this quantity and cannot evaluate).

Additionally, participants were asked to identify any items that were missing from the initial list of items, which they deemed to be important. The goal of this round was to obtain quantitative insights into the importance of specific concepts and gather additional items that should have been included in the preliminary work. The threshold for definite inclusion in Round 2 was that ≥75% of all responses (including those that responded “X – Don’t know or recognize this quantity and cannot evaluate”) needed to be above the midpoint of the scale (i.e. >4). Items with ≥75% of responses below the midpoint (<4) were excluded. Items which did not fall into these threshold-based categories were reserved for further evaluation in Round 2 and face-to-face discussions in Round 3. At the conclusion of Round 1, an initial draft checklist was made based on the ratings from experts for items within the predefined categories.

#### Round 2: Draft checklist evaluation and format specification

In Round 2, the draft checklist was circulated to experts for initial feedback. We asked experts to indicate whether there were any items from Round 1 which were not included in the draft that they feel should have been and whether there were any items included in the draft which should not have been. Additionally, we asked experts to indicate the preferred format (text, table figure) for each of the items included in the draft checklist. New items identified in Round 1 were introduced in Round 2 and expert consensus was sought for their inclusion. A draft checklist and results of Round 2 were then circulated to participants, along with an invitation to join the face-to-face discussions in Round 3.

#### Round 3: Face-to-face feedback and discussion sessions

In Round 3, the steering committee led 1-hour session and discussion rounds with small groups of participants via Zoom video calls. Only participants who completed both Round 1 and Round 2 were invited to participate. At least two members of the steering committee attended each session: one chairing the session and one taking notes. The sessions were also recorded so that they could be later reviewed if necessary. The sessions involved semi-structured discussions around the following themes: (1) clarifying any open questions or concerns; (2) discussing the scope of the guidelines and accompanying checklist; and (3) discussing dissemination and impact of the checklist and guidelines. Following the discussion sessions in Round 3, the checklist was revised to incorporate the feedback of the expert panel, and the accompanying guidelines for completing the checklist were written.

#### Round 4: Essential reporting items and guidelines

In Round 4, we sought consensus on which items on the checklist should be deemed essential to report regardless of context. That is, all items that were rated by more than 75% of the participants were considered essential and the “Not applicable” option was removed for these items in the final checklist. In addition, we also requested qualitative feedback on the final checklist draft and accompanying guidelines. Lastly, we evaluated the satisfaction of our participants with the process, and the resulting checklist and guidelines. Participants who completed both Round 1 and Round 2 (but not necessarily Round 3) were invited to participate.

Following Round 4, the ENLIGHT Steering Committee prepared the final checklist and guidelines, which were distributed to participants who accepted the invitation to join the ENLIGHT Consortium.

## Supporting information

ENLIGHT Checklist

ENLIGHT Guidelines

## Acknowledgements

We thank all participants of the consensus exercise, not all of whom completed all rounds and/or agreed to be named co-authors in the group membership model. This work was facilitated by participation in the NetIAS (Network of European Institutes for Advanced Study) Constructive Advanced Thinking (CAT) programme. We would like to thank Zukunftskolleg Konstanz, Aarhus Institute of Advanced Institute and Wissenschaftskolleg Berlin for organizing our visits. We are grateful for Uta Benner coordinating our participation in the NetIAS CAT programme.

## Author contributions

*Conceptualization:* M.S., L.K., R.L., E.M., R.P.N.

*Data curation:* n/a

*Formal Analysis:* n/a

*Funding acquisition:* M.S., L.K., R.L., E.M., R.P.N.

*Investigation:* M.S., L.K., R.L., E.M., R.P.N.

*Methodology:* M.S., L.K., R.L., E.M., R.P.N.

*Project administration*: n/a

*Resources*: n/a

*Software:* n/a

*Supervision*: n/a

*Validation*: n/a

*Visualization:* M.S., L.K., R.L., E.M., R.P.N.

*Writing – original draft:* M.S., L.K., R.L., E.M., R.P.N.

*Writing – review & editing:* M.S., L.K., R.L., E.M., R.P.N.

## Author order

The author order was decided as follows. M.S. is the lead of the NetIAS Constructive Advanced Thinking group project. The author order of L.K., R.L., E.M. and R.P.N. is alphabetical by last name. All authors contributed to the project equally.

