## Supplemental Materials for "ENLIGHT Consensus Checklist and Guidelines for reporting laboratory studies on the non-visual effects of light in humans"


Manuel Spitschan [0000-0002-8572-9268][1, 2, 3, +, *]

Laura Kervezee [0000-0002-6062-9164][4, +, *]

Renske Lok [0000-0003-1684-5625][5, +, *]

Elise McGlashan [0000-0002-3864-7198][6, +, *]

Raymond P. Najjar [0000-0002-3770-2300][7, 8, 9, 10, +, *]

for the ENLIGHT Consortium[¶]

[1] TUM School of Medicine & Health, Department of Health and Sport Sciences, Technical University of Munich, Munich, Germany

[2] TUM Institute for Advanced Study (TUM-IAS), Technical University of Munich, Garching, Germany

[3] Max Planck Institute for Biological Cybernetics, Max Planck Research Group Translational Sensory & Circadian Neuroscience, Tübingen, Germany

[4] Laboratory for Neurophysiology, Department of Cellular and Chemical Biology, Leiden University Medical Center, Leiden, Netherlands

[5] Department of Psychiatry and Behavioral Sciences, Stanford University, Stanford, USA

[6] School of Psychological Science and Turner Institute for Brain and Mental Health, Monash University, Melbourne, Australia

[7] Department of Ophthalmology, National University of Singapore, Singapore, Singapore

[8] Center for Innovation & Precision Eye Health, National University of Singapore, Singapore, Singapore

[9] Singapore Eye Research Institute, Singapore, Singapore

[10] Ophthalmology and Visual Sciences Academic Clinical Programme, Duke-NUS Medical School, Singapore, Singapore

+ Equal contribution


¶ The full list of ENLIGHT Consortium members can be found in **Supplemental Table S1** of this manuscript.

### Supplemental Material

**Supplemental Figure S1:** Ratings of inclusion and exclusion of items in the initial draft checklist (Round 2).

**Supplemental Figure S2:** Format preference of items in the initial draft checklist (Round 2).

**Supplemental Figure S3:** Percentage of votes for essential reporting of items (Round 4).

**Supplemental Table S1:** ENLIGHT Consortium Members.

**Supplemental Table S2:** Summary of reported light metrics in 19 studies described in Brown et al. (2020).

**Supplemental Table S3:** Common clusters of feedback raised in face to face discussion sessions in Round 3.

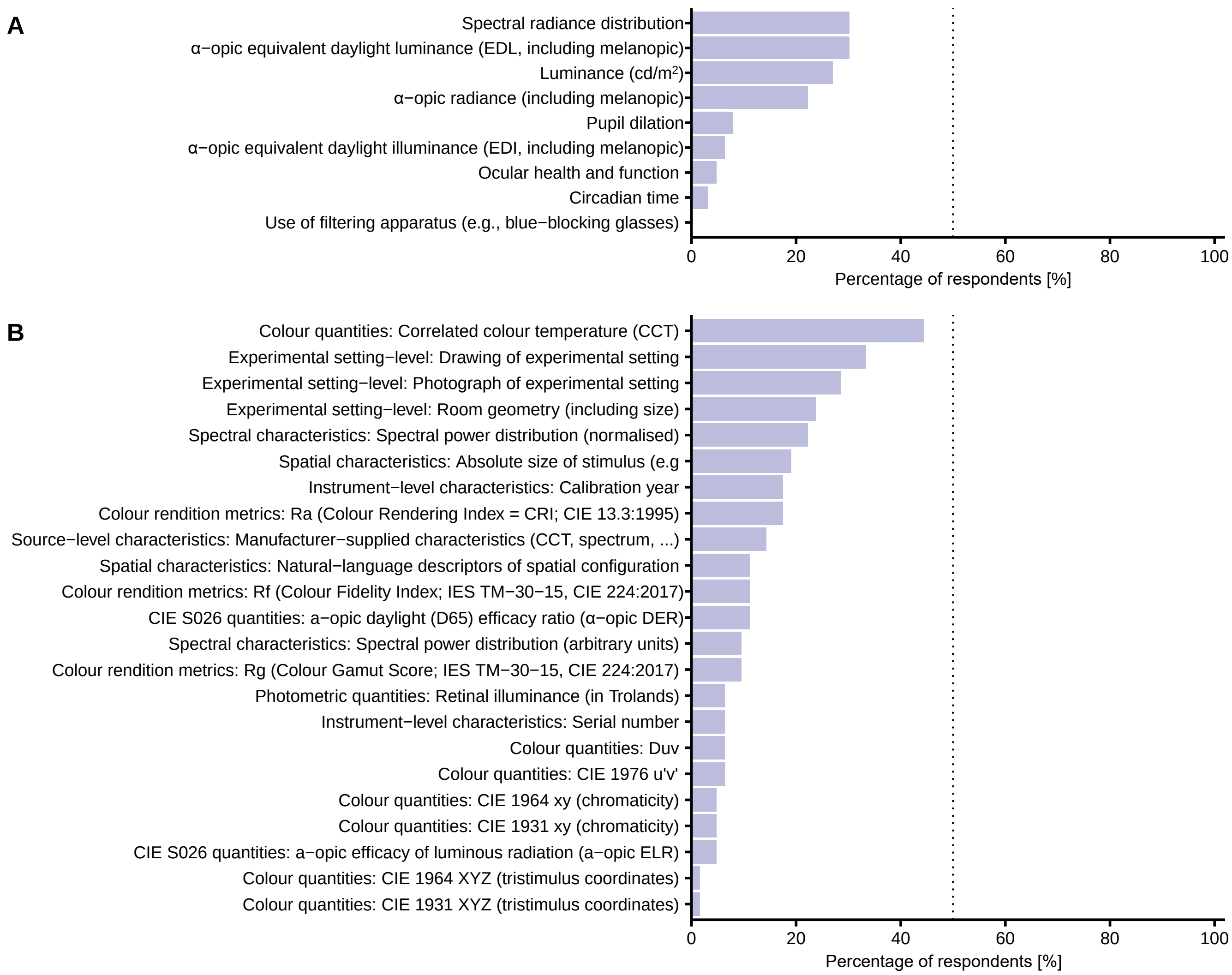


**Supplemental Figure S1:** Ratings of inclusion and exclusion of items in the initial draft checklist in Round 2 (n=63 participants). (A) Percentage of participants that rated a post hoc-included item as "should not have been included". (B) Percentage of participants that rated an excluded item as "should have been included".

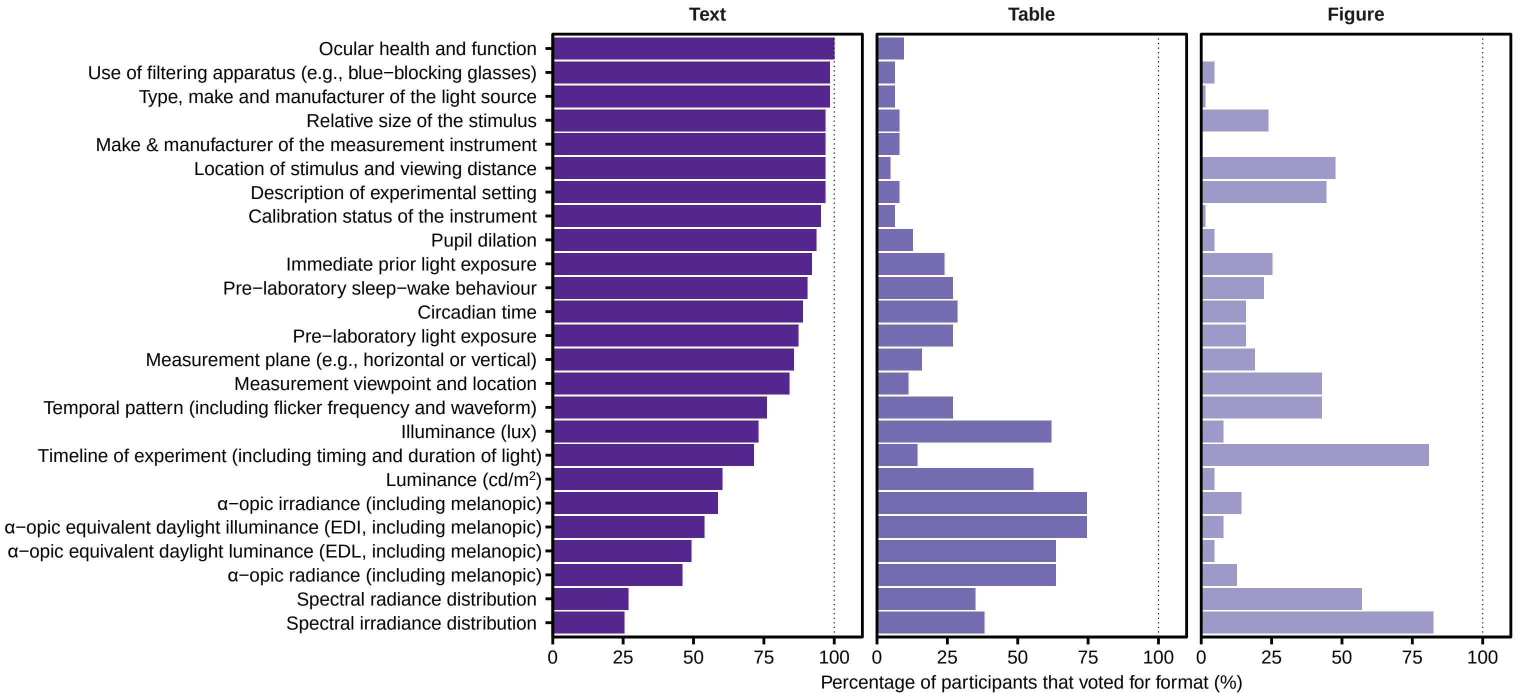


**Supplemental Figure S2:** Format preference of items in the initial draft checklist as rated by the participants (n=63 participants) in Round 2.

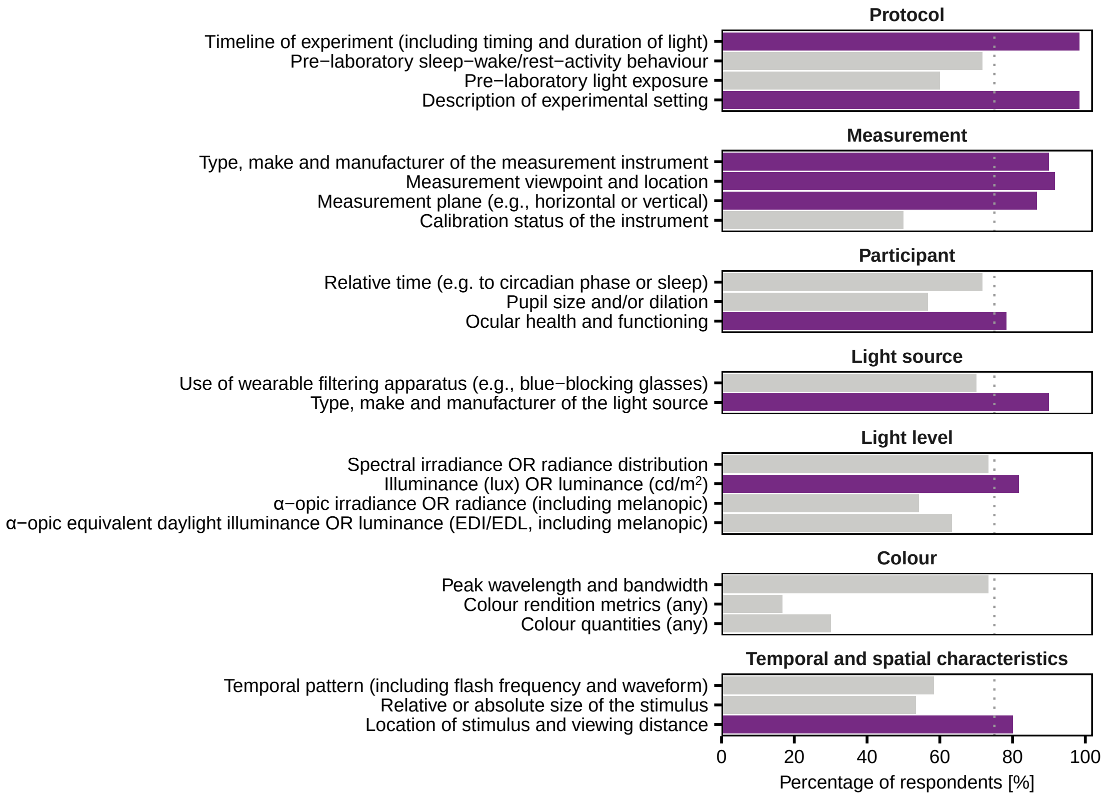


**Supplemental Figure S3:** Percentage of votes for mandatory reporting of items regardless of experimental context (n=60 participants in Round 4). Dark purple bars: items that reached consensus on being essential (i.e., were considered essential by more than 75% of respondents). Light gray bars: items that were voted by at most 75% of respondents as essential and therefore did not reach consensus on being essential to report regardless of context.

**Supplemental Table S1:** ENLIGHT Consortium members.

| Last name | First name | ORCID |
|---|---|---|
| Allen | Annette E. | 0000-0003-0214-7076 |
| Andersen | Marilyne | 0000-0001-8813-1184 |
| Bará | Salvador | 0000-0003-1274-8043 |
| Blattner | Peter | 0000-0003-0681-2096 |
| Blume | Christine | 0000-0003-2328-9612 |
| Boivin | Diane B. | 0000-0002-3896-2710 |
| Bonmatí-Carrión | María-Ángeles | 0000-0002-9050-6448 |
| Broszio | Kai | 0000-0002-8269-8654 |
| Brown | Timothy M. | 0000-0002-5625-4750 |
| Chellappa | Sarah Laxhmi | 0000-0002-6190-464X |
| Duffy | Jeanne F. | 0000-0003-4177-4179 |
| Eto | Taisuke | 0000-0002-0347-7408 |
| Flynn-Evans | Erin | |
| Fotios | Steve | 0000-0002-2410-7641 |
| Gabel | Virginie | 0000-0002-9792-1468 |
| Garbazza | Corrado | 0000-0002-8606-2944 |
| Glickman | Gena | 0000-0001-7064-7238 |
| Gordijn | Marijke C. | 0000-0001-9521-8085 |
| Hanifin | John P. | |
| Hartstein | Lauren | 0000-0002-3643-8894 |
| Herf | Michael | |
| Higuchi | Shigekazu | 0000-0001-7131-0792 |
| Hilditch | Cassie J. | 0000-0001-8797-4390 |
| Houser | Kevin W. | 0000-0001-6097-1560 |
| Hurlbert | Anya | 0000-0002-9879-5758 |
| LeBourgeois | Monique K. | |
| Lockley | Steven | 0000-0001-5209-2881 |
| Lucas | Robert | 0000-0002-1088-8029 |

*- Supplemental table S1 continues on next page -*

*- Supplemental table S1 continued from previous page -*

| | | |
|---|---|---|
| Moreno | Claudia R. C. | 0000-0003-1839-9673 |
| Münch | Mirjam | 0000-0003-2087-9916 |
| Mure | Ludovic S. | 0000-0002-2485-1032 |
| Peirson | Stuart | 0000-0003-3653-834X |
| Rahman | Shadab | 0000-0002-8729-0500 |
| Revell | Victoria L. | 0000-0002-8809-4587 |
| Rodriguez | Roberto G. | 0000-0001-7888-2152 |
| Roecklein | Kathryn | 0000-0001-5206-8283 |
| Rukmini | A. V. | 0000-0002-1300-7442 |
| Sammarco | John | 0000-0001-5843-634X |
| Santhi | Nayantara | |
| Schlangen | Luc J. M. | 0000-0002-8424-7240 |
| Schöllhorn | Isabel | 0000-0001-9067-1614 |
| Sharkey | Katherine M. | 0000-0003-1743-0704 |
| Skene | Debra J. | 0000-0001-8202-6180 |
| Sletten | Tracey L. | 0000-0002-0005-7838 |
| Smolders | Karin C. H. J. | 0000-0001-5838-8501 |
| Stefani | Oliver | 0000-0003-0199-6500 |
| Stone | Julia E. | 0000-0001-7706-7471 |
| Teikari | Petteri | 0000-0003-1095-4185 |
| Terman | Michael | 0000-0002-8871-3018 |
| Tran Quoc | Khanh | |
| Tsubota | Kazuo | 0000-0002-8874-7111 |
| Udovicic | Ljiljana | 0000-0002-6035-8309 |
| Vandewalle | Gilles | 0000-0003-2483-2752 |
| Veitch | Jennifer A. | 0000-0003-3183-4537 |
| Vetter | Céline | 0000-0002-3752-1067 |
| Wu | Lisa M. | 0000-0003-2241-9905 |
| Zauner | Johannes | 0000-0003-2171-4566 |
| Zeitzer | Jamie | 0000-0001-6174-5282 |

**Supplemental Table S2.** Summary of reported light metrics of 19 laboratory studies investigating effects of light on melatonin suppression and/or circadian phase resetting in response to at least two spectrally distinct sources delivered at multiple intensities. All articles were scored for reporting measures of light intensity, spectrum, and spectral power distribution.

| Category | Output measure | No. of studies reporting (out of 19) |
|---|---|---|
| Absolute spectrally unweighted intensities | Irradiance (energy units) | 10 |
| | Irradiance (photon units) | 11 |
| | Radiance (energy units) | 0 |
| | Radiance (photon units) | 0 |
| Light intensity | Illuminance (lux) | 14 |
| | Luminance (cd/m$^2$) | 1 |
| | Luminous flux (lumens) | 0 |
| | Any other luminance-like quantity | 1 |
| | Melanopic quantity | 8 |
| | Chromaticity (x,y, chromaticity coordinates) | 0 |
| | Correlated Colour Temperature | 8 |
| | Type of spectrum used | 19 (monochromatic: 10) |
| Narrowband spectra | Peak wavelength given | 10 |
| | Bandwidth given | 10 |
| Spectral power distribution | Absolute spectrum given (tabulated) | 2 |
| | Relative spectrum given (tabulated) | 0 |
| | Absolute spectrum given (graph) | 3 |
| | Relative spectrum given (graph) | 9 |

**Supplemental Table S3.** Summary of the most recurring feedback and suggestions, and actions taken by the steering committee following the synchronous discussions of Round 3.

| | **Feedback from ENLIGHT Consortium** | **Actions taken by the ENLIGHT Steering Committee** |
|---|---|---|
| **Objective 1:** To clarify open questions and concerns from participants | The checklist should be concise | To ensure conciseness, the following items were removed:<br>- Items that describe similar or alternative metrics.<br>- Preferred format (table, text, figure). |
| | Avoid the checklist being considered prescriptive or used to evaluate study quality. | It was clarified in the guidelines and manuscript that it is not the intention of the checklist to evaluate study quality. The checklist is meant to provide a space for authors to indicate whether and where metrics are described. |
| | Clarify whether NA means “not applicable” or “not available”. | Both “not applicable”, and “not available” were added to the checklist, as the committee appreciated that both responses may be relevant across items and it is important to provide the authors the opportunity to indicate either. |
| | Some light metrics may not be reportable by labs with limited resources. | As highlighted in the guidelines, most of the items included on the checklist can be measured and reported with minimal resources to ensure the checklist is as accessible as possible. |
| | The rationale regarding items included post-hoc following Round 1 should be made clearer to avoid subjectivity bias. | Following Round 1, none of the items included in the checklist were deemed by participants to be “unimportant” or “should not be included”. However, some aspects of lighting where *no individual items* reached the threshold for *definite inclusion (i.e., 75% consensus)*. Additional items that fell minimally below the threshold were added to the checklist, to ensure all aspects of light were covered. In Round 2, we asked participants to indicate whether they felt that post hoc included items should *not have been included*. For each of these items, less than 31% of experts rated them as “should not have been included”, and therefore all were retained. |

| | | |
|---|---|---|
| **Objective 2:** To discuss the scope of the guidelines accompanying the checklist | The guidelines should also highlight why it is important to take the light measurements specified in the checklist. | The description of why metrics or items are important is beyond the scope of the guidelines and checklist. The goal of this study is to reach consensus among experts on items to be reported in manuscript in order to enhance reproducibility and external validity of studies in the research field. |
| | The guidelines should mention that the objectives of the checklist are to enhance reproducibility and external validity of findings. | The objectives of the guidelines were revised to reaffirm the rationale and intention of the checklist and guidelines. |
| | The guidelines should address that some metrics are redundant and can be converted from other metrics. | Resources for converting metrics are provided in the guidelines, and this has been highlighted in the manuscript. |
| | The guidelines could include examples to indicate what quantity is important to report in what experiment, and which are critical/strongly recommended or optional. | The consensus around which items should be reported regardless of context (i.e., to remove the ‘not applicable’ checkbox for those items”) was achieved in Round 4.<br><br>In addition, we have also clarified in the checklist and guidelines the contexts in which the use of irradiance/illuminance versus radiance/luminance metrics may apply. |
| | The guidelines should include sufficient background information so that it is clear to researchers who are new to the field. | In the guidelines, we have explained the concepts included in the checklist as clearly and concisely as possible, and provided additional links to materials which may be of use. |
| **Objective 3:** To discuss on how to maximize impact and facilitate the adoption of the ENLIGHT Checklist and Guidelines | Several participants across sessions agreed that the checklist, guidelines, and any publications resulting from this work should be open access to ensure its widespread availability. They suggested that endorsement by organizations, research groups, and conferences could increase visibility, and that social media should be used to promote it. In addition, they emphasized the importance of gaining the support of journal editors, as crossover between communities tends to be limited, and a snowball effect could be achieved by following the manuscript with multiple editorials. | |
