## Supplementary material for "ENLIGHT Consensus Checklist and Guidelines for reporting laboratory studies on the non-visual effects of light in humans": ENLIGHT Checklist

Release 1.0.0, 17 February 2023

### ENLIGHT Reporting Checklist

Below is the **ENLIGHT Reporting Checklist** for reporting ocular light exposures in human laboratory-based studies. We will strongly encourage that this checklist be used in conjunction with the **ENLIGHT Reporting Guidelines**. This checklist is intended both to help authors, reviewers, and editors in evaluating the completeness of reporting in submitted studies, and for documentation of studies after publication. In the location column, please indicate the page, figure, or table number where the item or description can be found. If an item is not available, please select "**Not available**". If you consider an item not to be applicable in your specific study design after consulting the guidelines, please select "**Not applicable**". Items which do not have the option to select "Not applicable" were rated by experts as applicable for all studies, regardless of context. If you are unable to provide the information, please select "Not available".

The **ENLIGHT Reporting Checklist** (this document) and the **ENLIGHT Reporting Guidelines** are released under the CC-BY-NC-ND License. For more information, please visit http://enlight-statement.org/.

#### General Information

**Author names:**

**Title of manuscript:**

**Date:**

#### A. Study Characteristics

##### A.1. Protocol-level characteristics

| | Location (page, figure, table number) | Not available | Not applicable |
|---|---|---|---|
| Description of experimental setting | | ☐ | |
| Timeline of experiment (including timing and duration of light) | | ☐ | |
| Pre-laboratory sleep-wake/rest-activity behaviour | | ☐ | ☐ |
| Pre-laboratory light exposure | | ☐ | ☐ |
| Immediate prior light exposure (in laboratory) | | ☐ | ☐ |

##### A.2. Measurement-level characteristics

| | | | |
|---|---|---|---|
| Measurement plane (e.g., horizontal or vertical) | | ☐ | |
| Measurement viewpoint and location | | ☐ | |
| Type, make and manufacturer of the measurement instrument | | ☐ | |
| Calibration status of the instrument | | ☐ | ☐ |

##### A.3. Participant-level characteristics

| | | | |
|---|---|---|---|
| Ocular health and functioning | | ☐ | |
| Pupil size and/or dilation | | ☐ | ☐ |
| Relative time (e.g. to circadian phase or sleep) | | ☐ | ☐ |

### B. Light characteristics

#### B.1. Light source type(s). Please select all that are relevant.

| Room illumination (overhead or other) | Emissive surfaces including displays (incl. light therapy devices) | Wearable light emitting glasses | Ganzfeld exposure | Other: |
|---|---|---|---|---|
| ☐ | ☐ | ☐ | ☐ | ☐ |

| Polychromatic light | Monochromatic or narrowband light |
|---|---|
| ☐ | ☐ |

| | Location (page, figure, table number) | Not available | Not applicable |
|---|---|---|---|
| Type, make and manufacturer of the light source | | ☐ | |
| Use of wearable filtering apparatus (e.g., blue-blocking glasses) | | ☐ | ☐ |

#### B.2. Light level characteristics

| | | | |
|---|---|---|---|
| Illuminance (lux) and/or luminance (cd/m$^2$) | | ☐ | |
| Spectral irradiance and/or radiance distribution | | ☐ | ☐ |
| α−opic irradiance and/or radiance (including melanopic) | | ☐ | ☐ |
| α−opic equivalent daylight illuminance and/or luminance (EDI/EDL, including melanopic) | | ☐ | ☐ |

**NOTE:** luminance and radiance metrics (as opposed to illuminance and irradiance) are mainly relevant for emissive surfaces.

#### B.3. Colour characteristics

| | | | |
|---|---|---|---|
| Peak wavelength and bandwidth | | ☐ | ☐ |
| Colour appearance quantities (any) | | ☐ | ☐ |
| Colour rendering metrics (any) | | ☐ | ☐ |

**NOTE:** peak wavelength and bandwidth are most relevant for monochromatic or narrowband light sources.

#### B.4. Temporal and spatial characteristics

| | | | |
|---|---|---|---|
| Location of stimulus and viewing distance | | ☐ | |
| Temporal pattern (including flash frequency and waveform) | | ☐ | ☐ |
| Relative or absolute size of the stimulus | | ☐ | ☐ |

Reset form to default values

Print as PDF
