## Supplementary material for "ENLIGHT Consensus Checklist and Guidelines for reporting laboratory studies on the non-visual effects of light in humans": ENLIGHT Guidelines

### ENLIGHT Reporting Guidelines

#### Introduction

The **ENLIGHT Reporting Guidelines** are designed to guide authors in completing the accompanying checklist for their manuscript on laboratory-based studies of the non-visual effects of light. The **ENLIGHT Reporting Checklist** should be used as an addition to any other relevant reporting checklist, for example the CONSORT checklist. It is important to note that the guidelines are intended to instruct authors on the completion of the checklist, but not on the measurement or calculation of different metrics. Neither the checklist nor guidelines are intended to evaluate the quality of a study or prescribe specific research practices, and many of the items can be reported with minimal resources. They are intended to assist authors in reporting their conditions in a way which enables reproducibility and ensures external validity. Some metrics can be calculated from others, but authors are encouraged to provide the most complete representation of their experimental light condition(s). Resources to guide authors in calculating or describing certain metrics are provided below.

When completing the checklist, please indicate the page, figure, and/or table number where the information can be located in the manuscript. If any metric is not applicable (the item or metric is not relevant to the study), or not available (the item or metric was not collected or measured), please tick the appropriate box.

The ENLIGHT Reporting Checklist and the ENLIGHT Reporting Guidelines (this document) are released under the CC-BY-NC-ND License. For more information, please visit http://enlight-statement.org/.

#### A. Study Characteristics

##### A1. Protocol-level characteristics

*Description of experimental setting:* a description of the experimental setting, with as much detail as needed to reproduce the experiment. This includes, but is not restricted to the room geometry, colour of the walls, whether participants were ambulatory, seated, or supine, and whether the ambient temperature was controlled. A drawing or picture of the experimental setting may be included for additional clarity.

*Timeline of experiment (including timing and duration of light):* an overview of the timing of key study events/protocols, including clock-time if applicable. This includes the timing and duration of the light exposure, and may also include factors such as pre-exposure laboratory activities (e.g., tasks/procedures), and the timing of the measurement of outcome variables.

*Pre-laboratory sleep-wake/rest-activity behaviour:* a description of pre-laboratory sleep-wake or rest-activity behaviour. This includes but is not restricted to i) any inclusion/exclusion criteria that were in place regarding the participants' sleep timing or chronotype; ii) any instructions that the participants received regarding their sleep-wake or rest-activity behaviour prior to entering the laboratory (e.g. regarding sleep timing or regularity); and/or iii) any measurements that were done to characterise participants' sleep-wake or rest-activity behaviour prior to entering the laboratory (e.g. using actigraphy, sleep diaries, or at-home EEG/PSG). This could also include a description of any tools used to measure sleep or rest-activity cycles, as well as the duration across which the measurement took place. There are no recommendations for specific metrics which should be included, and *any* which are available in the manuscript may be reported here.

*Pre-laboratory light exposure:* a description of pre-laboratory light exposure including whether participants were given any instructions related to light-exposure, or whether this was controlled in any other way. This could also include a description of any tools used to estimate the light exposure, as well as the duration across which the measurement took place. There are no recommendations for specific metrics which should be included, and *any* which are available in the manuscript may be reported here.

*Immediate prior light exposure (in laboratory):* a description of the in-laboratory light conditions immediately prior to the experimental light exposure being reported in the manuscript. This is post-admit light exposure which may either describe general ambient lighting prior to an experimental condition, or a specific intervention in the protocol such as a dim-light condition or dark adaptation.

#### A.2. Measurement-level characteristics

*Measurement plane (e.g., horizontal or vertical):* a description of the plane that the light measurement(s) were performed in for each of the characteristics reported in Section B. If multiple measurements are reported, these details could be included for each one.

*Measurement viewpoint and location:* a description of the specific location at which the light sensor was placed during the measurements reported in Section B, including the direction of the sensor. If multiple measurements or experimental light conditions are reported, these details could be included for each one.

*Type, make and manufacturer of the measurement instrument:* a description of the instrument being used to take each measurement reported in Section B including the manufacturer, type, make and or model of the device.

Type: for example spectrophotometer, luxmeter, radiometer
Make: model number or make of the specific device
Manufacturer: company name and address

*Calibration status of the instrument:* a description of the calibration status of the light sensor that was used to take each measurement reported in Section B. This may include when, relative to the measurements, the device was calibrated and who performed the calibration.

##### A.3. Participant-level characteristics

*Pupil size and/or dilation:* a description of pupil size and/or whether pupils were dilated during the experimental protocol. This may include information on whether pupil size was measured during the experiment (and if so, how) and/or whether pupil size was controlled, for example through pharmacological agents (and if so, how).

*Relative time (e.g. to circadian phase or sleep):* a description of the time of the experimental light exposure relative to the participants' sleep or circadian timing. This may either mean that the experimental light exposure was intentionally timed relative to sleep or circadian timing on the day of the experimental exposure (or from a prior measurement), or that sleep and/or circadian time was measured and the relative time has been reported/recorded post-exposure.

*Ocular health and functioning:* any details on health and functioning of the participants' eyes. This may include inclusion/exclusion criteria that were used in the study regarding ocular health and functioning (for example, colour blindness, ocular diseases, ocular-motor dysfunction, etc.) as well as any information on how ocular health and functioning was assessed (for example within the study protocol by an ophthalmologist, through medical history, self-reported, etc.).

#### B. Light characteristics

Definitions and descriptions below have been adapted from the CIE E-International Lighting Vocabulary (E-ILV) and the α−opic toolboxes (Lucas et al., 2014, Schlangen et al., 2021). More information can be found below.

##### B.1. Light source type(s). Please select all that are relevant.

A description of the background and experimental light sources, types and locations used in the study, please tick all relevant options listed below. If a device (e.g., emissive surface or light emitting glasses) are being used in an illuminated room, please select both sources and provide details of the combined illumination as well as individual sources. This could include a description of any static filtering materials applied to the light source (for example neutral density filters).

Options are:

- *Room illumination (overhead or other):* a wall, roof, or floor-mounted light source resulting in room illumination. Examples include: ceiling lights and background light.
- *Emissive surfaces including displays (incl. light therapy devices):* light stimulus restricted to a surface. For example: Screens, light therapy devices, stand-alone LED panels. For these types of devices, radiance and luminance-related metrics are the recommended metrics to characterise the light level (see Section B2).
- *Wearable light emitting devices:* wearable devices with the capability of acting as a light source. For example: light emitting therapy glasses.
- *Ganzfeld exposure:* light exposure with a uniform full-field.
- *Other:* other potential light delivery sources not specified under Section **B1**.

Please also indicate which of the following best describes your light source/exposure:

- *Polychromatic light:* light source with a broadband spectral distribution.
- *Monochromatic or narrowband light:* a monochromatic light source is characterised by a light around single wavelength, while narrowband light covers a narrow range of wavelengths or that have a small fractional bandwidth. The source would have a clearly defined peak wavelength with a defined full width at half max amplitude.

*Type, make and manufacturer of the light source:* a description of the type, make and manufacturer of the light source(s) used for the stimulation.

Type: Fluorescent T5 tubes, LED light bulb, LED panel, LED strips, etc.
Make: model number
Manufacturer: Company name and address

*Use of wearable filtering apparatus:* a description of any wearable device(s) that modifies the absolute flux level (radiant or luminous) or relative spectral distribution, or both, of light passing through it. For example, blue-blocking glasses. This may include a description of the make and model, as well as an indication of the impact on the spectral quality and/or amount of light.

#### B.2. Light level

This section includes metrics to describe the light level used in the study (e.g. during the intervention or immediately prior to starting the experiment). Definitions for each are provided below. Additionally, further resources and tools (where available) for calculating metrics are provided below.

*Irradiance:* the density of incident radiant flux with respect to area at a point on a real or imaginary surface. Irradiance is usually expressed in watt per square metre ($W \cdot m^{-2}$).

*Radiance*: the density of radiant intensity with respect to projected area in a specified direction at a specified point on a real or imaginary surface. Radiance is generally expressed in watt per square metre per steradian ($W \cdot m^{-2} \cdot sr-1$).

*Spectral irradiance and/or radiance distribution:* spectral irradiance distribution is the density of irradiance with respect to wavelength. Spectral radiance distribution is the density of radiance with respect to wavelength.

*Illuminance (lux) and/or luminance (cd/m2):* illuminance describes the density of incident luminous flux with respect to area at a point on a real or imaginary surface. Luminance describes the density of luminous intensity with respect to the projected area in a specified direction at a specified point on a real or imaginary surface.

*α−opic irradiance and/or radiance (including melanopic):* α−opic irradiance or radiance describes the irradiance of the five photoreceptors in the human eye, i.e., rods, S- M- L-cones, and melanopsin, calculated from the spectral irradiance or radiance of a light source using the α-opic action spectrum weighted spectral radiance or radiance.

α−opic equivalent daylight illuminance and/or luminance (EDI/EDL, including melanopic):
EDI describes the illuminance level of daylight (D65) producing an equal α−opic irradiance.
EDL describes the amount of daylight (D65) needed to achieve a given α-opic luminance.

#### B.3. Colour characteristics

*Peak wavelength and bandwidth:* The peak wavelength describes the wavelength at which the spectral distribution reaches its largest value. The bandwidth describes the range or width of the spectrum, and may be reported as either the range, or the full-width half-maximum (i.e., the width of the spectral peak at 50% of the maximum power). These metrics are primarily applicable for monochromatic and narrowband light sources.

*Colour appearance quantities (any):* This includes any metric describing position in a chromaticity diagram or color space, such as (x,y), L*a*b*, Jab, or correlated colour temperature, CCT (Tc)."

*Colour rendering metrics (any):* Any metric describing the effect of a light source on the perceived colour of objects by conscious or subconscious comparison with their perceived colour under a reference light source. For example, the Colour Fidelity Index, Rf.

##### B.4. Temporal and spatial characteristics

*Temporal pattern (including flash frequency and waveform):* any description of a segment of signals that recurs frequently in the whole temporal signal sequence, for example, flash frequency or inter-stimulus interval, square wave or sinusoidal waveform.

*Location of stimulus and viewing distance:* any information describing the location of the light stimulus used in the study relative to the participant, and/or the relative distance between the light stimulus used in the study and the participant.

*Relative or absolute size of the stimulus:* any metrics describing the size and/or shape of the light stimulus, or the relative size (in relation to the visual field) used in the study.

#### Supporting materials

***Official CIE documents***

CIE. (2018). *CIE S 026/E:2018: CIE System for Metrology of Optical Radiation for ipRGC-Influenced Responses to Light*. CIE Central Bureau. https://doi.org/10.25039/S026.2018

CIE. (2020). *CIE S 017/E:2020: ILV: International Lighting Vocabulary, 2nd Edition*. CIE Central Bureau.

CIE. (2022). *e-ILV*. Retrieved 2 February 2023 from https://cie.co.at/e-ilv

***Software to calculate α-opic quantities from spectra***

CIE. (2020). *α-opic Toolbox for implementing CIE S 026* [Software]. CIE Central Bureau. https://doi.org/10.25039/S026.2018.TB

CIE. (2020). *User Guide to the α-opic Toolbox for implementing CIE S 026*. CIE Central Bureau. https://doi.org/10.25039/S026.2018.UG

Spitschan, M., Nam, S., & Veitch, J. A. (2022). *luox: Platform for calculating quantities related to light and lighting* [Software]. Available from https://luox.app/.

Spitschan, M., Mead, J., Roos, C., Lowis, C., Griffiths, B., Mucur, P., Herf, M., Nam, S., & Veitch, J. A. (2021). luox: validated reference open-access and open-source web platform for calculating and sharing physiologically relevant quantities for light and lighting. *Wellcome Open Res*, *6*, 69. https://doi.org/10.12688/wellcomeopenres.16595.3

***Previous work on standardising reporting conditions***

CIE. (2020). *CIE TN 011:2020: What to document and report in studies of ipRGC-influenced responses to light*. CIE Central Bureau. https://doi.org/10.25039/TN.011.2020

Knoop, M., Broszio, K., Diakite, A., Liedtke, C., Niedling, M., Rothert, I., Rudawski, F., & Weber, N. (2019). Methods to describe and measure lighting conditions in experiments on non-Image-forming aspects. *Leukos*, *15*(2-3), 163-179. https://doi.org/10.1080/15502724.2018.1518716

Spitschan, M., Stefani, O., Blattner, P., Gronfier, C., Lockley, S. W., & Lucas, R. J. (2019). How to report light exposure in human chronobiology and sleep research experiments. *Clocks Sleep*, *1*(3), 280-289. https://doi.org/10.3390/clockssleep1030024

***General background on radiometry and photometry***

DeCusatis, C., & Optical Society of America. (1997). *Handbook of applied photometry*. AIP Press ;
Optical Society of America.

Price, L. L. A., & Blattner, P. (2022). Circadian and visual photometry. *Prog Brain Res*, *273*(1), 1-11. https://doi.org/10.1016/bs.pbr.2022.02.014

Schlangen, L. J. M., & Price, L. L. A. (2021). The lighting environment, its metrology, and non-visual responses. *Front Neurol*, *12*, 624861. https://doi.org/10.3389/fneur.2021.624861

Sliney, D. H. (2007). Radiometric quantities and units used in photobiology and photochemistry: recommendations of the Commission Internationale de L'Eclairage (International Commission on Illumination). *Photochem Photobiol*, *83*(2), 425-432. https://doi.org/10.1562/2006-11-14-RA-1081
